# NeuroSuite for Long-term Functional and Structural Studies of Air-Liquid Interface Cerebral Organoids

**DOI:** 10.1101/2025.07.16.663353

**Authors:** Belquis Haider, Sagnik Middya, Daniel Lloyd-Davies-Sánchez, Nino Laubli, Sulay Vora, Jakob Träuble, Rohan Krajeski, Rubén Ruiz-Mateos Serrano, Ole Paulsen, Madeline Lancaster, George G. Malliaras, Gabriele Kaminski Schierle

## Abstract

Over the past decade, air-liquid interface cerebral organoids (ALI-COs) have emerged as powerful in vitro models that capture essential structural and functional traits of the human brain, offering an exciting alternative to traditional animal models in neuroscience. Yet, the full potential of these systems has remained untapped due to the lack of non-invasive, long-term electrophysiological tools capable of preserving organoid integrity. Existing techniques, ranging from patch clamping to rigid and 3D microelectrode arrays, often compromise organoid growth and disrupt delicate cytoarchitecture. Here, we present NeuroSuite, an innovative bioelectronic platform designed to overcome these challenges. At its core is Neuroweb, a perforated, ultra-thin, and conformable organic microelectrode array engineered for minimal disruption of nutrient and oxygen exchange. Neuroweb is reusable and supports stable recordings for over six months, making it uniquely suited for longitudinal studies. Coated with poly(3,4-ethylenedioxythiophene):polystyrene sulfonate (PEDOT:PSS), a high-performance mixed ionic-electronic conductor, Neuroweb delivers exceptional signal-to-noise ratio recordings with high spatial precision. By pairing Neuroweb with NeuroMaps, an intuitive software for interactive analysis and visualisation, NeuroSuite enables long-term, non-invasive tracking and spatial mapping of electrical activity from brain organoids and ex vivo brain slices at the air-liquid interface. Following rigorous validation, we demonstrate that NeuroSuite can capture both high- and low-frequency throughout maturation. Our pipeline reveals evolving network connectivity, including the development of GABA-ergic interneurons, and concurrent shifts in high-frequency spiking and low-frequency oscillations indicative of a refinement in the excitatory-inhibitory balance. Finally, automated data acquisition and spatial spike mapping highlight local activity changes in response to media composition, a factor often overlooked in conventional recordings. NeuroSuite thus opens a new frontier in organoid neuroscience, enabling precise, long-term monitoring essential for modelling neurological diseases, understanding human brain development, and accelerating drug discovery.

**Teaser:** Conformal organic bioelectronic arrays, combined with an open-access toolbox for analysis and visualisation, reveal real-time and long-term neural dynamics in brain organoid slices at the air-liquid interface.

## Introduction

Microelectrode arrays (MEAs) have become essential tools for recording the electrical activity of biological tissues, enabling insights into cellular dynamics and network behaviour at high-resolution. Originally developed to overcome the low throughput and invasiveness of techniques like patch clamping in 2D cultures^1^, MEAs have evolved in tandem with advances in tissue modelling, most notably, the emergence of 3D brain organoids. Derived from human embryonic stem cells, brain organoids are self-organising, three-dimensional structures that recapitulate key aspects of human brain morphology and function^2,3^. They offer unprecedented opportunities for studying neurodevelopment^2,4–6^, neurodegeneration^7–9^, and neurotoxicity^10,11^, while circumventing many of the ethical and translational limitations of animal models ^12^.

However, as organoids grow in size and complexity, they face challenges in oxygen and nutrient diffusion that can hinder maturation and viability^13^. To address this, air-liquid interface cerebral organoids (ALI-COs) have been developed, enhancing neuronal survival and axonal outgrowth by exposing one surface to air while maintaining contact with the culture medium ^14,15^. Despite these innovations, the long-term, *in situ* electrophysiological monitoring of brain organoids remains a significant technical hurdle^16 17^. Traditional MEAs, whether building on rigid planar surfaces or emerging 3D and mesh-like constructs, often compromise the organoid’s structural integrity, interfere with tissue development, or require re-plating steps that disrupt cytoarchitecture^6,18–21^. Moreover, the mechanical mismatch between stiff electrode materials and soft brain tissue can affect the neuronal natural organisation or alter signalling, further confounding results^22–24^.

Recent advances in bioelectronics have led to the development of flexible and even stretchable MEAs using materials with lower Young’s moduli than glass, including SU-8, parylene-C, styrene-ethylene-butylene-styrene (SEBS), and polydimethylsiloxane (PDMS), and with form factors that better accommodate tissue mechanics^25–28^. These have improved interfacing in cardiac cells and neural tissue *in vivo*, but their adaptation for brain organoid research is still emerging. One state-of-the-art approach introduced mesh nanoelectronics that integrate into brain organoids during early development, enabling chronic, single-cell recordings^28^. However, this method requires embedding during morphogenesis and is limited to 3D organoids that often develop necrotic cores, thus restricting long-term culture while being incompatibility with slicing protocols due to the risk of damaging to the embedded device. Other strategies, such as kirigami-inspired^29^ or shell-type^27^ mesh arrays, offer improved experimental flexibility and mechanical adaptability, but are often hindered by complex assembly and handling procedures. Moreover, their geometries typically span ~1 mm from the organoid base, making them incompatible with ALI-COs that expand to diameters of ~5 mm^32–34^.

To overcome these limitations, we introduce Neuroweb, a bioelectronic interface tailored for long-term, non-invasive monitoring of brain organoids and ex vivo brain slices cultured at the air-liquid interface. Neuroweb features a perforated, ultra-thin (4 μm) parylene-C substrate with a low 30% fill ratio, offering excellent conformability while preserving nutrient and gas exchange. The device’s 32 gold (Au) electrodes are coated with poly(3,4-ethylenedioxythiophene):polystyrene sulfonate (PEDOT:PSS), yielding a high signal-to-noise ratio (~16 dB) and spatial resolution (~30 μm), ideal for capturing fine-scale neural activity as slices mature from 500 μm to 5 mm.

We validate Neuroweb through comprehensive functional characterisation and demonstrate its performance in long-term electrophysiological recordings of cortical ALI-COs. By combining it with NeuroMaps, our open-access, interactive software suite for data analysis and visualisation, we tracked organoid electrical activity, quantifying and mapping key metrics such as impedance, along with spike rates, synchrony, bursting patterns, and waveform features (e.g., full-width half-maximum, amplitude, skewness). The impedance remained stable for up to six months while the waveform features evolved dynamically with organoid maturation and were indicative of emerging neural connectivity. Crucially, NeuroMaps revealed a progressive shift in spike population activity which was likely driven by the maturation of the neuronal population and increasing inhibitory activity that regulated neuronal patterns. This was accompanied by changes in the aperiodic exponent of the local field potential (LFP) power spectrum, an indicator of network excitation-inhibition balance^30^, alongside increased burst regularity and spatially localised connectivity, allowing detailed assessments of network topology. Together, Neuroweb and NeuroMaps establish a powerful, scalable framework for long-term functional studies of human brain organoids, revealing a level of insight that would be inaccessible using conventional, short-term devices or platforms which decouple high- and low-frequency analyses, thus paving the way for new insights in disease modelling, developmental neuroscience, and high-throughput drug screening.

## Results

NeuroSuite has been specifically designed for non-invasive, long-term electrophysiological monitoring of ALI brain organoids. The NeuroSuite workflow is depicted in **Figure 1a**, in which organoid slices are generated independently of Neuroweb’s fabrication. Arrays are fabricated using scalable microfabrication techniques and arrays are integrated into a customisable, autoclavable polycarbonate enclosure compatible with standard cell-culture inserts. This setup supports stable organoid maintenance and routine media exchange, while minimising evaporation and contamination. Thus, the system is fully compatible with long-term electrophysiology, imaging, and perfusion experiments.

**Figure 1.**
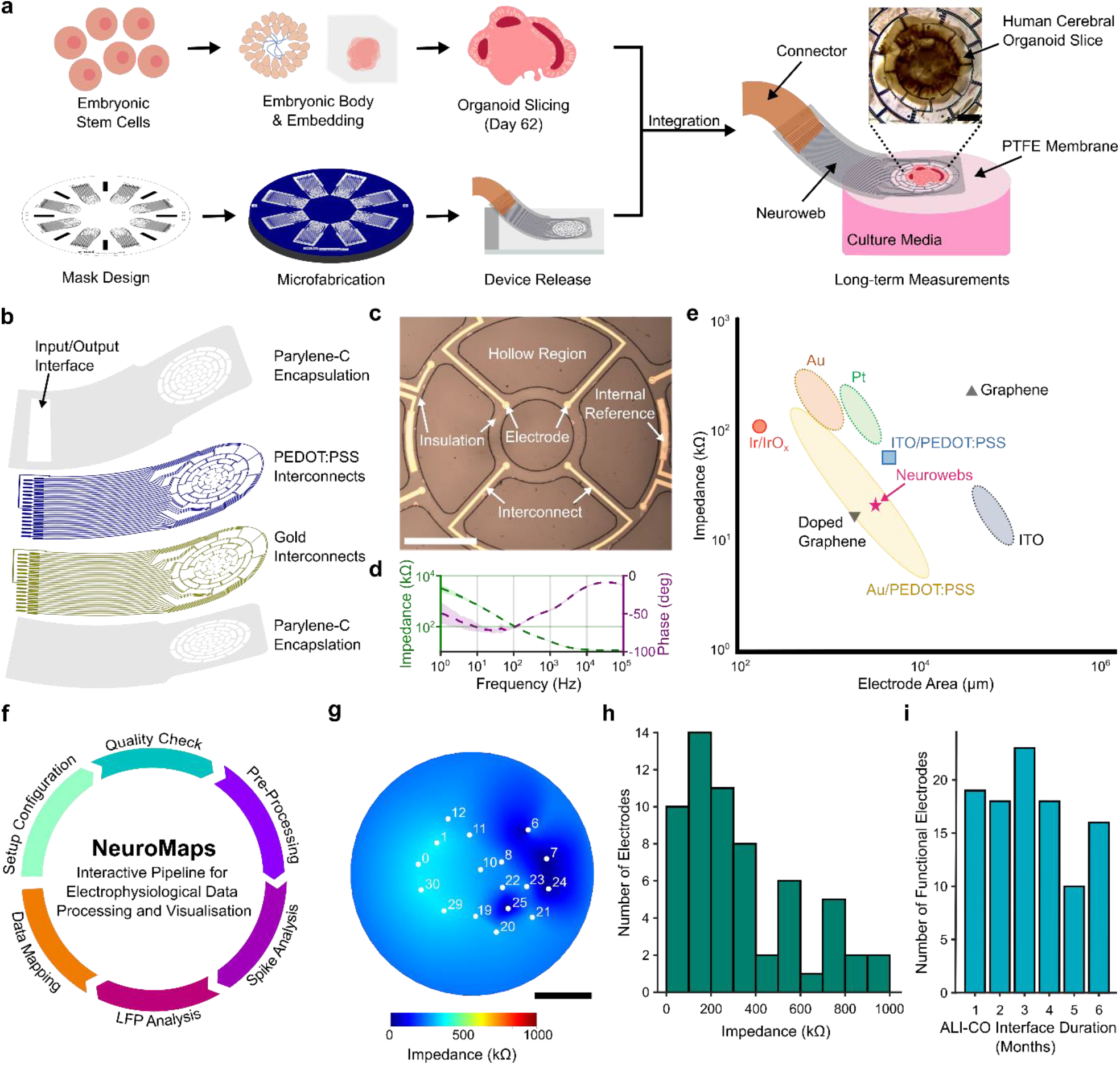
The NeuroSuite pipeline: device design, fabrication, and functional characterisation for long-term integration with ALICOs. (a) Overview of the parallel workflow showing the generation of cerebral organoid slices from human embryonic stem cells, microfabrication of Neuroweb devices, and their integration for long-term culture at the air-liquid interface. (b) Zoomed-in schematic of Neuroweb’s multilayer structure: a 2 μm top Parylene-C encapsulation layer, a 600 nm PEDOT:PSS electrode layer, a 100 nm gold interconnect layer, and a 2 μm bottom Parylene-C substrate. (c) Optical image of Neuroweb highlighting the perforated, ultrathin structure that enables efficient nutrient exchange and conformal interfacing with organoid slices; and scanning electron microscopy (SEM) images Error! Reference source not found.. (d) Bode plot showing impedance and phase profiles of PEDOT:PSS-coated gold electrodes (1 Hz-100 kHz), reflecting typical capacitive behaviour at low frequencies and resistive behaviour at higher frequencies. (e) Benchmark comparison of Neuroweb’s electrode impedance (at 1 kHz in PBS) with various materials reported in the literature, including Au^32–34^, Au/Pt^29,33^, Pt^35,36^, Ir/IrOx ^37^, Si ^37^, ITO^38,39^, and ITO/PEDOT:PSS^40^. (f) Illustration of NeuroMaps software functions, enabling comprehensive analysis of Neuroweb data including signal filtering, quality tracking, spike detection, spatial mapping, and extraction of electrophysiological features. (g) Impedance heatmap of all electrodes mapped onto the array layout using the NeuroMaps software platform. (h) Histogram of impedance values from two devices integrated with organoids, exhibiting a Gamma distribution and demonstrating that most electrodes remain below 200 kΩ at 1 kHz. (i) Histogram quantifying the number of functional electrodes across six months of continuous organoid interfacing, assessed via impedance, standard deviation, median absolute deviation, and power spectral density (see Materials and Methods). Scale bars: (a) 1 mm, (c) 500 μm, (g) 1 mm.

Neuroweb’s 4.4 mm electrode span accommodates organoids growing from ~700 μm at day 62, i.e., the day of slicing and integration, to ~5 mm at late stages. The device integrates 32 gold electrodes coated with PEDOT:PSS and encapsulated in a 4 μm-thick biostable Parylene-C film (**Figure 1b-c**). Electrodes are arranged in five concentric rings with 600 μm radial spacing and include internal references optimised for both multi-unit activity (MUA) and local field potential (LFP) recordings. Perforations across ~70% of the central area ensure unobstructed nutrient diffusion and reduce the risk of necrosis or cellular stress at the device-tissue interface. The array’s bending stiffness (~0.72 pNm^2^), estimated using classical beam theory and accounting for both geometrical and material contributions, is sufficiently low to minimise the mechanical mismatch with organoid slices at the air-liquid interface^24^. It also provides structural stability to ensure robust handling, interfacing, and long-term electrophysiological interrogation of soft tissue.

The PEDOT:PSS coating is a mixed ionic-electronic conductor^36^ that increases the capacitance of the electrode/electrolyte interface^31^ (**Figure 1d**). Combined with the underlying low-resistance gold layer ^21^, Neuroweb achieves a low impedance of 25 ± 9 kΩ at 1 kHz in phosphate-buffered saline (PBS) at 25 ºC (**Figure 1e**). Complementing Neuroweb, we introduce NeuroMaps, a MATLAB-based platform for electrode quality assessment, processing longitudinal electrophysiological data processing, and spatial mapping (**Figure 1f** Error! Reference source not found.). Using this software, we spatially mapped impedance values across the organoid (**Figure 1g**), revealing a Gamma-like distribution with most electrodes exhibiting an impedance of under 200 kΩ at 1kHz (**Figure 1h**). Notably, over 50% of electrodes remain functional after six months of continuous interface and multiple ethanol device clean-up cycles (**Figure 1i**).

### Neuroweb Supports Long-Term Growth and Maturation of ALICOs

To assess Neuroweb’s biocompatibility and suitability for long-term organoid interfacing, we cultured ALI-COs directly on the device. Cortical organoids were generated from human embryonic stem cells which were seeded, differentiated, and embedded in Matrigel following established protocols ^14^. Slicing using microtomy was carried out at 59-62 days in vitro (DIV) to preserve structural integrity before transfer onto the Neuroweb platform. Cultures were maintained at the ALI for an additional 48 days (up to DIV 110) as described in the Materials and Methods section, after which samples were fixed and processed for imaging.

Immunofluorescence confirmed robust organoid expansion across the device, with no evidence of major necrosis (DAPI) or impaired growth (**Figure 2a**). Neuroweb’s perforated design and minimal insulating footprint permitted tissue integration, enabling dense networks of neurons (MAP2) and astrocytes (GFAP) to envelop the electrode architecture (**Figure 2a**, top inset). Axonal projections extended throughout the slice, and MAP2 staining revealed radially aligned dendrites, indicative of healthy, directional neuronal outgrowth (**Figure 2a**, bottom inset).

**Figure 2.**
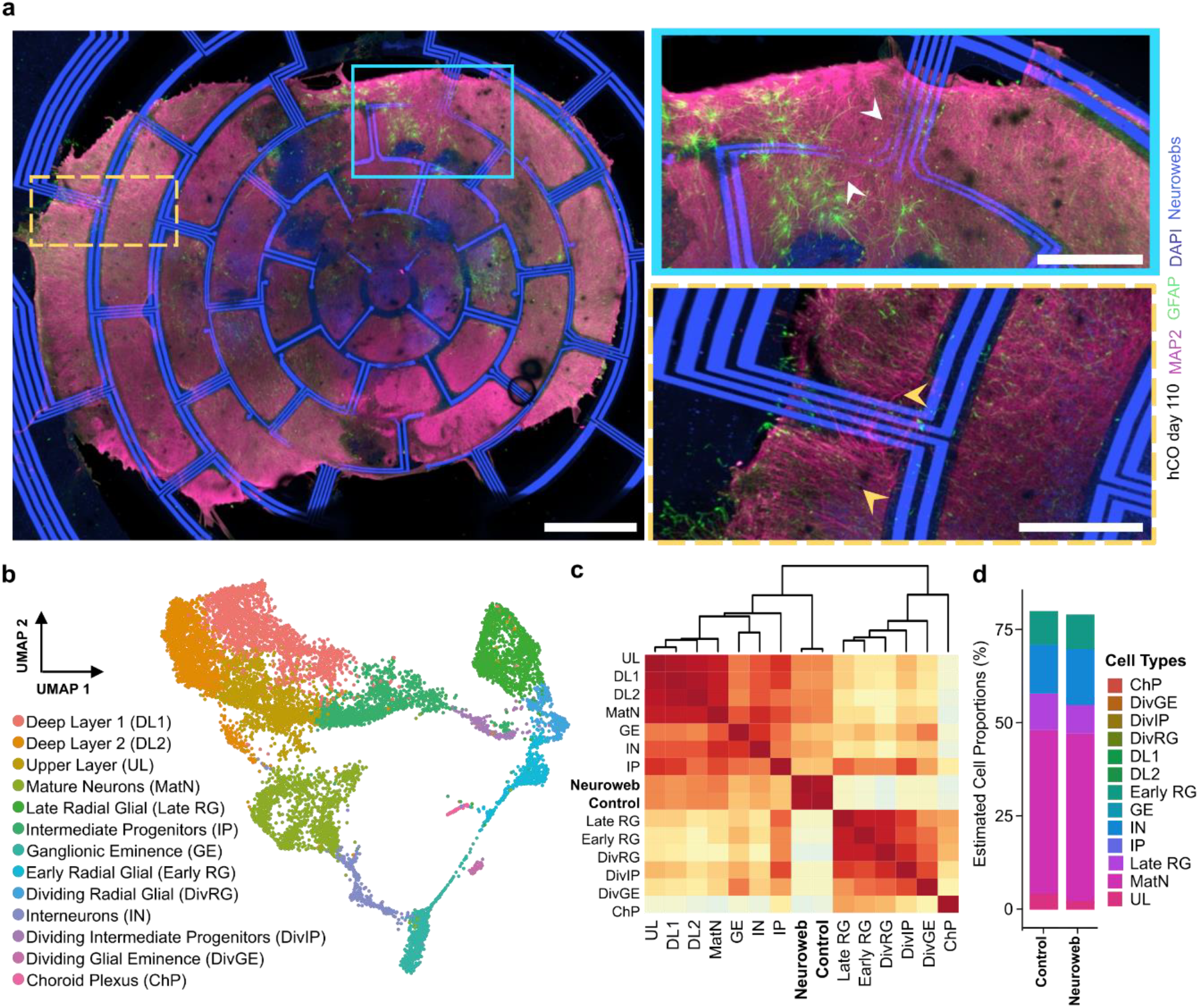
Neuroweb supports robust growth, structural organisation, and cellular diversity in ALI-COs. (a) Confocal immunofluorescence image of a human cerebral organoid cultured on Neuroweb for 48 days (DIV 110). Nuclei are stained with DAPI (dark blue), neurons with MAP2 (magenta), and glial cells with GFAP (green); the Neuroweb conductive tracts appear in bright blue. The top-right inset highlights organoid encapsulation of the device (white arrows), while the bottom-right inset shows radially aligned MAP2-positive dendrites (yellow arrows), indicative of directional neuronal outgrowth. (b) Uniform Manifold Approximation and Projection (UMAP) plot of single-cell RNA sequencing data from Neuroweb-supported organoids on DIV 110, identifying 13 major cell clusters. (c) Hierarchical clustering dendrogram showing dominant mature neuronal populations—upper-layer (UL), deep-layer 1 and 2 (DL1, DL2), mature neurons (MatN), ganglionic eminence (GE), interneurons (IN), and intermediate progenitors (IP)—with only minor representation of progenitor and non-neuronal types, including late, early, and dividing radial glia (LateRG, EarlyRG, DivRG), dividing IPs (DivIP), dividing GE cells (DivGE), and choroid plexus (ChP). (d) Bar plot showing cell-type proportions derived from deconvolved single-cell RNA-sequencing (scRNA-seq) data for Neuroweb-grown organoids and controls (cultured on standard cell-culture inserts), demonstrating comparable cellular composition. (c–d) n = 2 biological replicates per condition. Scale bars: (a) Main panel: 1 mm; insets: 400 μm.

To further evaluate the cellular composition of Neuroweb-supported organoids, we performed RNA sequencing and compared the results to single-cell data of standard ALI-COs. Uniform Manifold Approximation and Projection (UMAP) analysis identified thirteen major cell types, including upper and deep-layer excitatory neurons (UL, DL1, DL2), interneurons (IN), radial glia (RG), intermediate progenitors (IP), and choroid plexus cells (ChP) (**Figure 2b**). Crucially, Neuroweb-grown ALI-COs displayed cellular diversity comparable to controls cultured on standard inserts (**Figure 2c-d**) and exhibited similarity to more mature cell types similar to control (**Figure 2c**), confirming that the device does not perturb natural developmental trajectories.

Finally, to assess the significance of Neuroweb’s perforated architecture, we compared it to a micro-electrocorticographic array (μECoG), previously validated in vivo^41^, composed of the same materials but with a higher fill ratio^41^. After 69 days of co-culture, immunostaining revealed a stark contrast in biocompatibility. Neuroweb with a 30% fill ratio supported extensive neuronal (MAP2) and axonal (SMI312) networks alongside healthy glial morphology. In contrast, organoids cultured on the higher-density μECoG (70% fill ratio) exhibited minimal growth and large necrotic regions, likely due to impaired nutrient diffusion.

### NeuroSuite Enables Long-Term Electrophysiological Recording and Visualisation of ALI-CO Network Dynamics

Following validation of Neuroweb’s suitability for long-term culturing of ALI-COs, we developed NeuroMaps as a modular, interactive software toolbox designed for longitudinal analysis and visualisation of spontaneous and evoked electrophysiological activity. The pipeline includes integrated pre-processing, post-processing, and spatial mapping functionalities.

Pre-processing included impedance-based channel selection (<1 MΩ), noise thresholding, and the application of established quality metrics^42^ (detailed in Materials and Methods Error! Reference source not found.). Representative raw signals from selected channels, containing both low- and high-frequency components, are shown across developmental timepoints (**Figure 3a**). Early recordings (DIV 62) revealed sparse, uncoordinated neuronal spiking activity, which evolved into broader network-wide activity by DIV 82. Near-synchronous spike initiation across multiple electrodes at DIV 72 suggests increasing gap junction-mediated synchronisation. Meanwhile, slight temporal delays between signals on spatially separated electrodes indicate activity from distinct neuronal populations, validating Neuroweb’s capacity to resolve inter-population dynamics.

**Figure 3.**
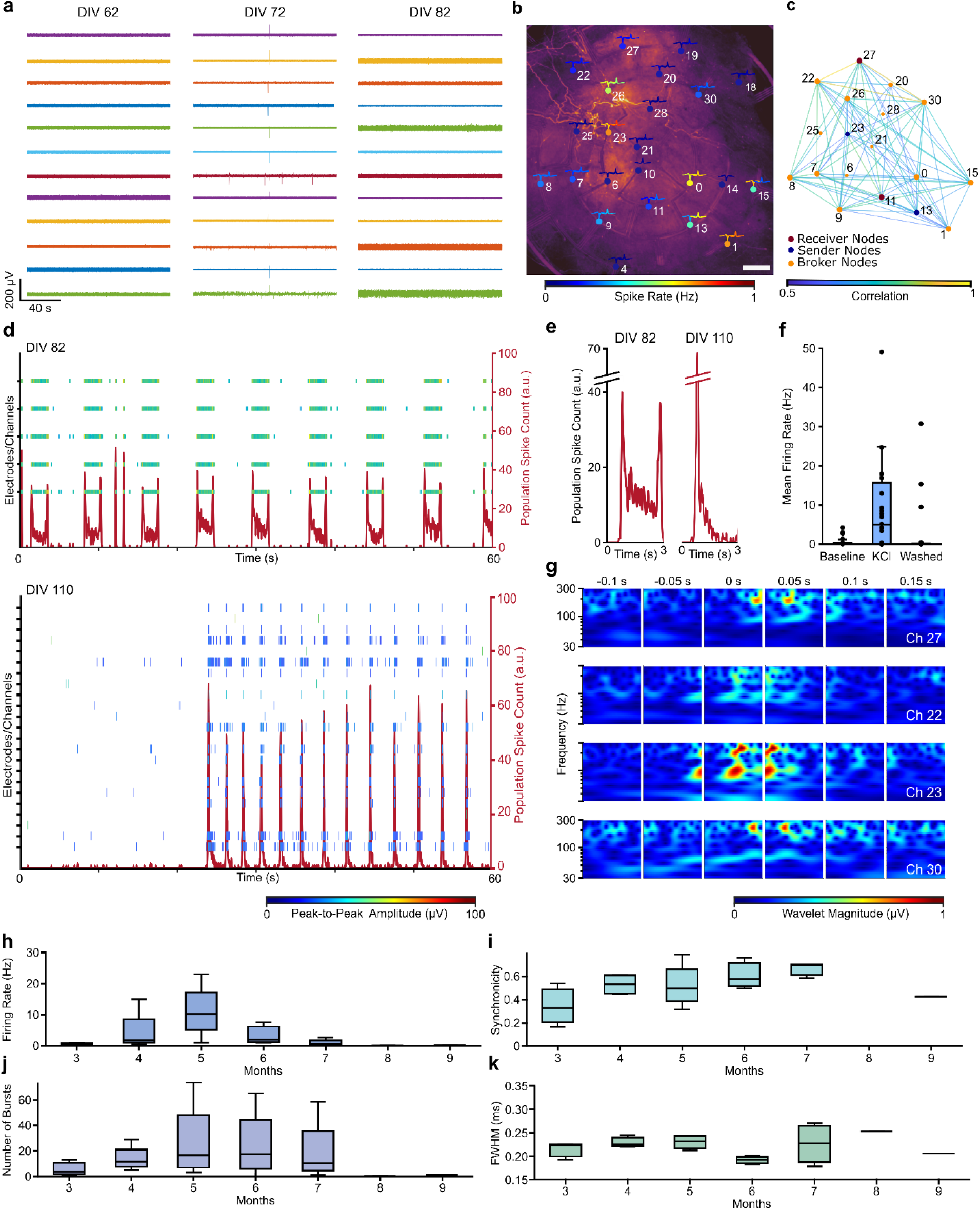
Longitudinal tracking of electrophysiological maturation in ALI-COs. (a) Representative raw traces of spontaneous activity recorded from an organoid slice at DIV 62, 72, and 82, illustrating progressive increases in network activity. (b) Detected action potentials overlaid on a corresponding immunofluorescence image using NeuroMaps, enabling spatial co-registration of electrical and anatomical data. (c) Functional connectivity map based on STTC analysis, identifying key network nodes; electrode 27 is classified as a receiver and electrode 22 as a broker. (d) Raster plots overlaid with population spike rate plots at DIV 82 and DIV 110 reveal increasing temporal precision and network synchrony during organoid maturation. The colour scale indicates spike peak-to-peak amplitude. (e) Zoomed-in traces of network-wide spiking events highlight changes in activity dynamics between DIV 82 and DIV 110, with sharper and more transient responses at later stages. (f) Validation of neuronal activity via stimulation with 50 mM KCl, which significantly increased spike activity across 11 channels (n = 2 organoids). (g) Continuous wavelet transform analysis showing broadband gamma bursts (30-300 Hz) during population activity. Temporal alignment of bursts across channels 27 and 22, as well as 30 and 23, corresponds to network roles inferred in (c). (h–k) Longitudinal analysis of key electrophysiological metrics: (h) mean firing rate, (i) network synchronicity, (j) spike FWHM, and (k) burst frequency. Firing rate peaks at month 5 before declining, while synchrony increases progressively until month 7. Scale bar: (b) 500 μm.

To quantify coordinated neuronal activity, spikes were extracted from band-pass filtered signals (300-3000 Hz) using an amplitude threshold of 4.5× the noise standard deviation^43^, a 2 ms refractory period, and a spike feature filter based on a full-width half maximum (FWHM) criterion of 1 ms to exclude artefacts. Using NeuroMaps, we localised spike origins at the electrode level and co-registered waveforms with immunofluorescence images (**Figure 3b**). Given the high cellular density, these signals reflect multi-unit or multi-source activity within densely packed networks.

To assess spatial and functional connectivity^44^, we computed the ‘spike-time tiling coefficient’ (STTC) matrix^45^ across all channel pairs, classifying electrodes as senders, receivers, or brokers based on the ratio of incoming to outgoing spikes^14,46^ (**Figure 3c**). This analysis revealed the strongest correlations at an interelectrode distance of 1359 μm, with moderate correlations peaking at 1738 μm and weaker correlations around 1639 μm, consistent with Neuroweb’s concentric design and supporting the presence of long-range, spatially organised network activity.

Raster plots and population activity traces (computation detailed in Materials and Methods) at DIV 82 and DIV 110 (**Figure 3d-e**) illustrated distinct maturation stages. At DIV 82, spontaneous activity was prolonged and variable, while at DIV 110, it became more transient with sharply decaying profiles. This shift likely reflects the maturation of inhibitory circuitry and more refined network excitability, driven by the expected increase in GABAergic interneurons during cortical maturation^47–49^. Biological signal origin was confirmed by depolarising two organoids with potassium chloride (KCl) ^50^, which significantly increased activity; signal levels returned to baseline upon washout (**Figure 3f**).

We next analysed low-frequency oscillatory dynamics using continuous wavelet transform (CWT) to capture both temporal and spectral characteristics of population activity (**Figure 3g**). This revealed co-occurring bursts in the broadband gamma range (30-300 Hz) during episodes of high-frequency neuronal spiking activity, oscillations associated with local circuitry spanning 250-500 μm in vivo ^51,52^. In organoids, these are likely driven by excitatory, inhibitory interactions shaping population-level output. Notably, a temporal lag between bursts on channels 22, 23, 27, and 30 aligned with their inferred functional roles from the STTC map: channel 23 as a sender, channel 27 as a receiver, and the others as brokers (**Figure 3c**). This behaviour was also similar in an independent measurement, whereby spike feature analysis depicted a change in the peak-to-peak amplitude of the spikes during propagation while the full-width-half-maximum remained relatively constant.

Beyond the snapshot analyses presented above, the high biocompatibility and long-term stability of Neuroweb enable NeuroSuite to extract and analyse longitudinal, high-frequency electrophysiological features from ALI-COs. These include core activity metrics such as mean spike rate, network synchrony, and burst frequency, as well as signal waveform properties like FWHM. These analyses were based on 5-minute-long recordings acquired from eleven cerebral organoids across partially overlapping time windows. Notably, one organoid remained continuously interfaced with Neuroweb for up to six months—without repositioning or manual adjustment—demonstrating the platform’s suitability for stable, chronic recordings.

The average spike rate increased from 0.87 ± 0.18 Hz at month 3 post-seeding to a peak of 11.2 ± 9 Hz at month 5, followed by a progressive decline to 3.6 ± 1 Hz in month 6 and 1.3 ± 1.04 Hz in month 7 (**Figure 3h**). Beyond this point, activity fell below 0.3 Hz, correlating with a significant drop in the number of neuronal activity (<10 electrodes detected activity). Relatedly, network synchronicity rose from 0.342 ± 0.19 to 0.52 ± 0.2 between months 3 and 5 (**Figure 3i**), while burst frequency increased from 5.8 ± 6.1 to 27.4 ± 31.8 bursts per five-minute interval (**Figure 3j**), both indicative of maturing connectivity and coordinated neuronal spiking activity. Interestingly, despite these functional changes, FWHM values remained relatively stable across all time points (**Figure 3k**), suggesting that the biophysical properties of spike waveforms were maintained throughout extended culture. This consistency underscores Neuroweb’s ability to provide a minimally invasive, stable interface for chronic recordings, preserving signal fidelity even as network-level activity evolves.

### NeuroSuite Reveals Age-Dependent Shifts in Low-Frequency Network Dynamics

To extend our electrophysiological characterisation beyond high-frequency spiking activity, we analysed low-frequency components of the recorded signals, which offer complementary insight into large-scale network behaviour. While high-frequency signals (300-3000 Hz) capture spatially localised action potentials, LFPs in the 0.5-300 Hz range reflect ensemble-level neuronal dynamics across broader spatial domains, providing valuable markers of network synchronisation, oscillatory coordination, and temporal structure.

Focusing on mature ALI-COs at DIV 110, we decomposed broadband recordings into low-frequency LFP components and compared them to their corresponding high-frequency traces (**Figure 4a**). A magnified view during a population burst (**Figure 4b**) revealed a precise temporal alignment between high-frequency spiking and low-frequency oscillations, indicating coordinated activity across multiple spatial and temporal scales.

**Figure 4.**
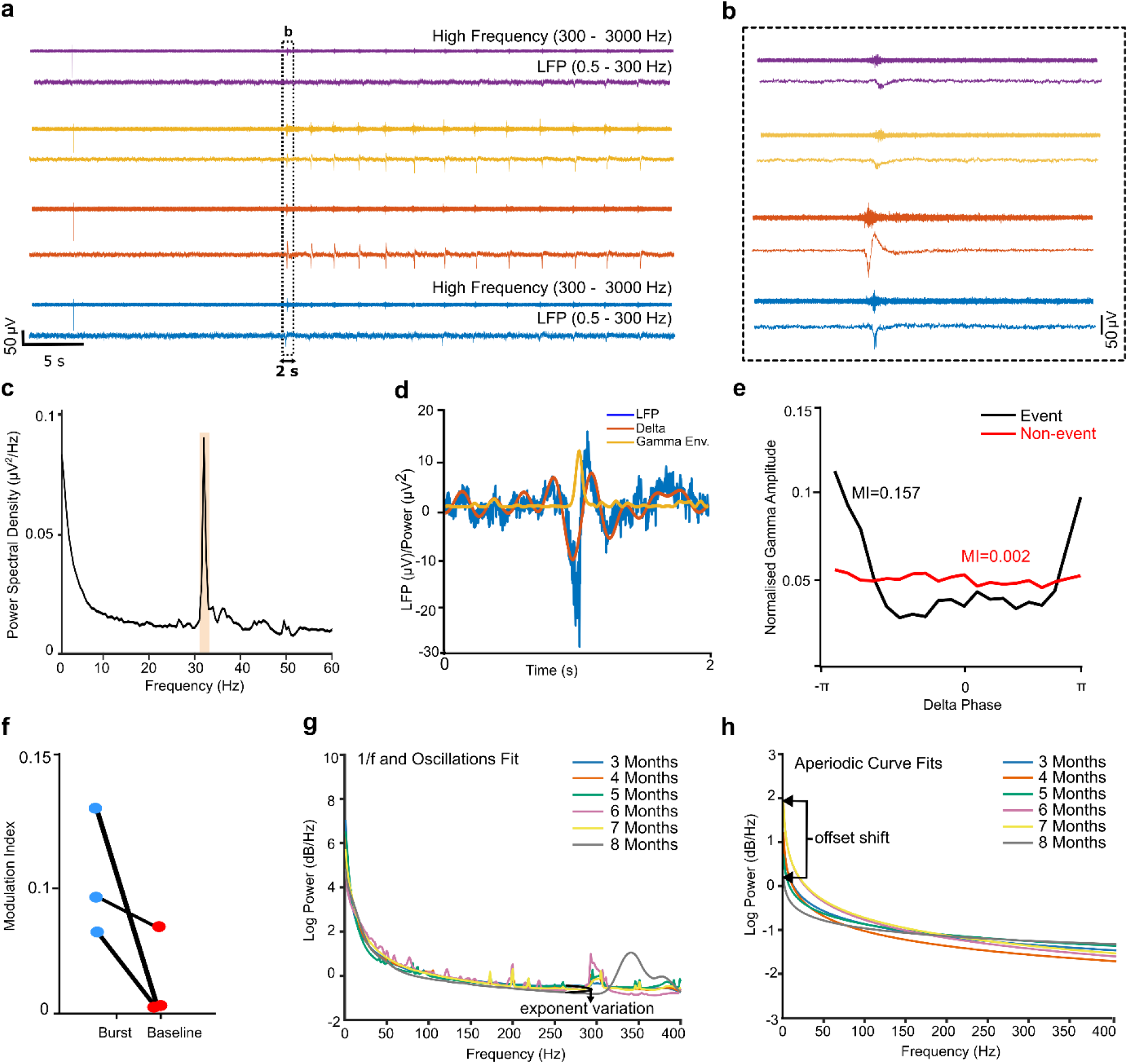
Spectral and temporal features of neural activity reveal coordinated network dynamics and maturation in ALI-COs. (a) Representative traces recorded on DIV 110 from multiple electrodes (each represented by a different color) show broadband (0.5–3000 Hz) and band-pass filtered signals. Top traces represent high-frequency activity (300– 3000 Hz), while bottom traces show low-frequency LFPs (0.5–300 Hz). (b) Zoom-in of a synchronised network burst reveals temporal alignment between spiking and LFP oscillations. (c) Power spectral density (PSD) from a representative mature organoid shows a peak in gamma activity around 35–40 Hz. (d) Decomposition of the LFP signal into broadband gamma amplitude (yellow) and delta phase (orange) illustrates cross-frequency modulation. (e) PAC analysis shows elevated modulation index (MI = 0.157, black) during network bursts compared to baseline periods (MI = 0.002, red). (f) Average PAC values confirm that gamma–delta coupling is enhanced during synchronised network events. (g–h) Parameterised PSD fits show age-dependent reductions in the aperiodic offset and exponent, consistent with neural network maturation. The arrow in (g) depicts the curves’ aperiodic (1/f) exponential variation over maturation.

To quantify these oscillations, we performed power spectral density (PSD) analysis of the LFP signals (**Figure 4c**). A distinct spectral peak emerged in the gamma range (30-60 Hz), which has been previously linked to synchronised population spiking activity^53^. Gamma power was notably elevated during bursts of spiking, reinforcing its value as a surrogate marker for coordinated ensemble activity.

Given the known interaction between fast (gamma) and slow (delta, 1-4 Hz) oscillations in cortical circuits, we examined phase-amplitude coupling (PAC) to determine whether high-frequency gamma activity was modulated by slower delta rhythms^6,54–56^. Signals were filtered using a finite impulse response (FIR) filter to extract gamma amplitude and delta phase (**Figure 4d**). The modulation index (MI), a commonly used metric for quantifying PAC, was significantly elevated during synchronised network events compared to baseline periods (MI: 0.157 vs. 0.002, **Figure 4e-f**). This suggests that fast dynamics are temporally structured by slow-wave activity, a hallmark of hierarchical organisation and functional maturation in both in vivo and in vitro neural systems^57,58^.

We further explored changes in aperiodic and periodic spectral features using spectral parameterisation algorithms^30^ that were embedded into NeuroMaps. In cortical ALI-COs, both the aperiodic exponent and offset increased up to month 7, suggesting a progressive development of the ALI-COs inhibitory circuits and enhanced population-level connectivity.

These trends may reflect the maturation of GABAergic signalling or a refinement in the excitatory-inhibitory balance^30^ (**Figure 4g-h**).

Lastly, we applied this algorithm to track oscillatory dynamics over 24-hour recordings, observing not only a maturation-associated increase in gamma power but also fluctuations in spectral power at ~14-hours of recording. This may indicate intrinsic metabolic cycling within the organoids, highlighting their potential as in vitro models for long-term neural metabolic and rhythm research.

## Discussion

In this work, we developed NeuroSuite, a next-generation platform that enables stable, high-resolution electrophysiological recordings from ALI-COs for over six months. At its heart is Neuroweb, a flexible, perforated organic microelectrode array that supports the dynamic growth of sliced organoids (up to 5 mm diameter), while preserving metabolic exchange and tissue integrity. Paired with NeuroMaps, a user-friendly MATLAB-based toolbox for interactive analysis and visualisation, our platform enables researchers to capture, visualise, and quantify evolving high-and low-frequency neural dynamics over extended timescales.

While traditional rigid and penetrating microelectrode arrays have been used in organoid research^6,14^, offering valuable functional insights through their high-throughput capabilities^46,59^ and improved signal-to-noise ratios^60^, they often fall short in chronic applications due to their rigidity, invasiveness, and tendency to disrupt tissue architecture^22,23^. Neuroweb overcomes these limitations through its ultra-thin, low-fill design, which promotes nutrient and oxygen diffusion without compromising spatial resolution. Unlike SU-8 substrates used in flexible mesh arrays^28,29^, and which has been linked to cytotoxicity^61,62^, parylene-C supports spontaneous growth and encapsulation of ALI-COs. Immunofluorescence imaging and RNA-sequencing confirmed biocompatibility, enabling a stable tissue-electrode interface for high-fidelity signal acquisition from air-liquid interface cultures.

Moreover, Neuroweb integrates seamlessly with standard cell-culture components through a modular, 3D-printable, and autoclavable holder compatible with standard inserts and imaging systems. This setup enables *in situ* recordings within incubators, reducing handling complexity^26,29^ and eliminating the need of manual gluing steps for device stabilisation and assembly^63^, while supporting spontaneous tissue development. PEDOT:PSS-coated gold electrodes further elevate Neuroweb’s performance, achieving superior signal-to-noise ratios relative to platinum^28,35,36^, ITO^38,39^, uncoated gold^29,32,33^, and graphene^35,64^, without compromising spatial resolution (30 μm).

NeuroSuite is the first platform to integrate minimally invasive, chronic interfacing with a user-friendly analytical suite – unlocking scalable, high-throughput interrogation of organoid network dynamics over time. NeuroMaps offers an intuitive graphical user interface built on validated standards^6,34,45,65–67^. Users can access a suite of tools for signal quality assessment, spike and burst detection, network mapping, low-frequency aperiodic, periodic, and oscillatory analysis with no prior coding experience necessary. Over six months of continuous interface, NeuroSuite enabled robust tracking of network activity in maturing ALI-COs, capturing dynamic changes in spike frequency, bursting patterns, and synchronicity. Beyond fast spiking, the platform revealed shifts in low-frequency oscillatory behaviour, including gamma–delta coupling and a rhythmic modulation in spectral power at the 14-hour mark after media exchange, pointing to intrinsic maturation and metabolic cycles.

Future extensions of Neuroweb, including larger stimulation electrodes^68,69^, biosensors^25^, or micro-light-emitting diodes (*μ*-LEDs)^70^ could enable multimodal circuit probing and neuromodulation. Additionally, integrating scalable data handling frameworks and machine learning-based spike sorting into NeuroMaps would enable high-throughput studies across larger and more complex organoid models. Together, these innovations establish NeuroSuite as a transformative platform for probing and decoding circuit-level function, human brain development, and neurodegenerative disease progression. With its seamless integration of hardware and analysis, NeuroSuite paves the way for a new era of non-invasive, high-resolution, and scalable exploration of brain organoid function, advancing research not only in neurodevelopment but also in healthy ageing, age-related brain disorders, and pharmaceutical drug screening.

## Materials and Methods

### Flexible Device Design and Fabrication

The devices were designed using Clewin Layout editor and ordered as film photomasks (JD Photo Data). Microfabrication was carried out on a 4-inch silicon wafer which was sonicated sequentially in acetone followed by isopropanol, and dried thoroughly on a hot plate. A 2 μm layer of parylene-C was deposited onto the silicon wafer via chemical vapour deposition (CVD, Speciality Coating Systems Labcoater, USA). An image-reversal photoresist (AZ 5214E, Microchemicals GmbH, Germany) was patterned to define the electrode and interconnect geometry through photolithography (SUSS MABA6, Mask Aligner) and developed in an aqueous dilution of the developer AZ351B at a ratio of 1:4 (Microchemicals GmbH, Germany). This was followed by electron beam evaporation of the metal electrodes and interconnects (Ti, 10-nm adhesion layer and Au, 100 nm; PVD 75, Kurt J Lesker) and acetone lift-off. The PEDOT:PSS solution was prepared by mixing a PEDOT:PSS suspension (Clevios PH 1000, Heraeus, Germany), 5% (v/v) ethylene glycol, and approximately 30 μL dodecyl benzene sulfonic acid (Sigma Aldrich, UK). A cross-linker, 1% (v/v) 3-Glycidyloxypropyltrimethoxysilane (GOPS, Sigma-Aldrich, UK), was added and filtered through a 0.45 μm pore-size poly(tetrafluoroethylene) filter (Sartorius). This mixture was used within 20 minutes, by spin-coating in 4 layers onto the silicon wafer to create a 600 nm thick layer. The sample was baked for 1-minute at 110°C between each coating step, and at 120°C for 1 hour after the four layers were coated. Overnight soaking in deionised water was done to remove excess PSS and low-molecular-weight compounds. The electrodes were further defined by photolithographic patterning the PEDOT:PSS using a positive photoresist (AZ5214E) developed in the aqueous dilution of the developer AZ351B at a ratio of 1:4 (Microchemicals GmbH, Germany), and reactive ion etching of PEDOT:PSS (CF4, 50 sccm, SF6, 20 sccm, 30 mTorr, 200W; PlasmaPro RIE 80, Oxford Instruments, UK). Lastly, the wafer’s surface was plasma activated and immersed in a self-assembled monolayer (SAM) of the adhesion promoter of 3% (v/v) methacryloxypropyl trimethoxysilane (A174 silane, Sigma Aldrich, UK) prepared in 96% ethanol (containing 1% acetic acid) for 30 seconds followed by an ethanol rinse and heat treatment on a hot plate at 70^0^C for 1 hour. This enhanced the adhesion of the subsequently deposited 2 *μm* parylene-C encapsulation layer (CVD, Speciality Coating Systems Labcoater, USA). Finally, two separate patterning steps were performed to define the device outline and the electrode openings, respectively, using a positive photoresist (AZ10XT) in conjunction with AZ 726 MIF. Etching for each patterning step was carried out using a reactive ion etcher (CF4, 8 sccm; SF6, 2 sccm; O2, 50 sccm; 60 mTorr) (PlasmaPro RIE 80 from Oxford Instruments, UK). The flexible devices were gently peeled off from the silicon wafer while soaked in deionized water, transferred onto glass slides using a paint brush, and dried on an 80°C hot plate.

### Device Release and Cable Bonding

Jumper cables (FFC/FPC, Molex 15014-0613) were bonded using anisotropic conductive adhesive film (ACF) (TGP5010 with 10 µm particle size, Telephus Inc., Korea) and a die bonder (FINEPLACER pico2, Finetech GmbH, Germany). The bonded probes were subsequently released from the glass slides using deionised water, and a low-toxicity fast-curing silicone adhesive (Kwik Sil™) was used to further seal the bonded interconnects area.

### Design and assembly of 3D printed culture platform

The 3D printed culture platform consists of three components, the air-liquid interface compatible insert, the holder, and the PCB plug-in module. These are designed to be compatible with the dish sizes used in this work but can be adapted according to demand. All 3D printed components are fabricated using polycarbonate filament and sterilized in three steps. First, all components are washed in soapy water, followed by 70% ethanol rinse, and U.V. sterilization for 20 minutes. These components are also compatible with autoclaving. The insert is first assembled with the desired tubing of the culture platform and air-liquid interface inserts was performed using OnShape. The culture platform was 3D printed using a filament printer and polycarbonate.

### Electrochemical Impedance Spectroscopy

Electrochemical Impedance Spectroscopy was carried out with a potentiostat (Autolab Potentiostat, Metrohm AG, Switzerland) using a platinum (Pt) electrode as the counter electrode and silver-silver chloride (Ag/AgCl) reference electrode. A 10-mV amplitude sinusoidal voltage input with frequencies ranging from 1 Hz to 100 kHz was used. The impedance of PEDOT:PSS was additionally measured using an Intan RHS stim/recording system (Intan Technologies, Los Angeles, CA) in 1x phosphate-buffered saline (PBS) at 1 kHz.

### Scanning Electron Microscope Imaging

The SEM images were taken using a ZEISS Gemini 300 VP scanning electron microscope using an acceleration voltage of 15 kV in backscattered electron (BSE) mode. The flexible device mounted on a microscopy slide was placed on top of carbon tape and analysed in point-by-point scanning mode.

### Human stem cell, organoid, and ALI-CO culture

H9 (human female) embryonic stem cells purchased from WiCell (WA09) were approved for use in this project by the UK Stem Cell Bank Steering Committee. Cells were maintained in StemFlex medium (ThermoFisher Scientific, A3349401) on plates coated with Matrigel (Corning, 356234) and passaged twice weekly with 0.1M EDTA. The commercially available STEMdiff Cerebral Organoid Kit (StemCell Technologies, 08570) was used to generate cerebral organoids, seeding 2000 cells per embryoid body. After 15 days in vitro (DIV), the cerebral organoids were excised from the Matrigel droplets in which they were embedded during the Kit protocol, with a needle and fine scalpel under a stereomicroscope and returned to organoid media. After DIV 30, organoid media was continually supplemented with Matrigel dissolved at 2% (v/v). For culture at the air-liquid interface (ALI), cerebral organoids (CO) of 55-70 (DIV) were processed to 300 μm slices as previously described^14^ and collected directly onto cell culture inserts (Millipore, PICMORG50) with serum free slice culture medium beneath (SFSCM: Neurobasal (ThermoFisher Scientific, 21103049), 1x B-27 supplement (ThermoFisher Scientific, 17504044), 1x Glutamax (ThermoFisher Scientific, 35050038), 0.5% (w/v) glucose). After two weeks of ALICO culture, the media was switched to BrainPhys Neuronal Medium with N2-A and SM1 (StemCell Technologies, 05793), supplemented with 35ng/ml vitamin C. All ALI-CO media were supplemented with 1x antibiotic-antimycotic (ThermoFisher Scientific, 15240062) and additional 1:1000 (v/v) fungizone (Merck, A2942), and media was refreshed daily. All cell, organoid, and ALICO cultures were maintained at 5% CO_2_ at 37 °C.

### ALI-CO culture on MEAs

The cell culture inserts, and associated Neuroweb were soaked in 70% ethanol for sterilisation and allowed to air dry. Before plating ALICOs on Neuroweb, the cell culture inserts, and Neuroweb were washed once in PBS and then three times in ALICO media. Neuroweb was placed centrally onto the surface of the cell culture inserts (Millipore, PICMORG50) and flattened using water and a fine paintbrush, so that surface tension stuck them to the surface. Once dry, ALI-CO slices were collected onto the cell culture inserts in the usual manner, and positioned so that they rested centrally on the Neuroweb in the centre of the insert. Gentle flooding of the top surface of the insert with ALICO media, so that ALICOs float, allowed manipulation and repositioning of the ALICO slice with a pipette tip to achieve optimal positioning before removal of the excess media on the insert.

### Immunofluorescence staining

ALI-CO fixation and immunohistochemistry ALI-COs were fixed in 4% PFA overnight at 4Cand subsequently washed three times in PBS. Staining of whole ALICOs was carried out as previously described^14^ with all steps performed in PBS with 0.25% Triton-X and 4% normal donkey serum. The following primary antibodies were used: GFAP (Abcam, ab7260, 1:1000 dilution), MAP2 (Abcam, ab5392, 1:200 dilution), SMI312 (Biolegend, #837904, 1:500 dilution). Secondary antibodies (Thermo Fisher Scientific and Abcam) with AlexaFluor 488-, 568-, and 647-labels were used for detection. Imaging of stained whole ALI-COs was performed using a Nikon CSU-W1 spinning disk microscope with a 10x air- or 25x silicon-immersion lens.

### RNA-sequencing (RNA-seq) and analysis

RNA extraction for all samples was conducted in parallel using the Direct-zol RNA extraction kit for 96 samples (Zymo Research, R2054), as per the manufacturer’s instruction. Poly(A) mRNA transcripts were isolated, and cDNA libraries were generated using NEBNext modules for library preparations (New England Biolabs, E7490 and E7760), as per the manufacturer’s instructions, and during which unique dual index UMI adapters (oligos for multiplexing sequencing) were incorporated (New England Biolabs, E7416 and E7874). Libraries were sequenced on a Nextseq 2000 with a P3 flow cell and cartridge. Bulk RNA sequencing data were aligned to genome assemblies using the NextFlow RNA-seq pipeline (version 3.12.0; nf-co.re/rnaseq/3.12.0/)^71^. Data were aligned to the Homo sapiens assembly GRCh38.p14 (GCA_000001405.29; release 102) and alignment employed the default RNA-seq pipeline parameters to output values expressed as transcripts per million (TPM).

Single cell data from ALICOs was from Giandomenico et al. (2019) and was reanalysed starting from the gene expression matrix using Seurat version 5.1.0 in R (version 4.4.0). Cells were filtered for healthy cells with less than 1.8% mitochondrial reads and nCount_RNA >2100 but less than 3000 and nFeature_RNA >800 but less than 1500 to exclude multiplets after exploring the nFeature_RNA x nCount_RNA graph. Data normalization, variable features and scaling were done according to the standard Seurat pipeline. Clustering was done with resolution set to 0.43 and UMAP visualization performed on the top 50 principle components. Clusters were annotated based on the top 20 differentially expressed genes according to log2FC, guided by key known marker genes. To compare with single-cell RNA-seq (scRNA-seq), the bulk RNA-seq analysis, the TPM matrix was first filtered for highly expressed genes with TPM greater than or equal to 1 and then log transformed for comparison to scRNA-seq. The clusters from scRNA-seq were used to generate pseudobulk data by first sub-setting the Seurat object by each cluster, followed by conversion to a sum gene expression matrix of the raw counts and then calculation of counts per million (CPM). Only genes with CPM of at least 1 in one sample were filtered, the data was then log transformed and merged with bulk TPM matrix. The top 2000 most variable genes were then taken forward visualization using Pearson correlation using heatmap.2 and PCA analysis using prcomp. Deconvolution was performed using the Bioconductor package, granulator version 1.12.0.

### Recording

A custom-made printed-circuit board was connected to the FFC/FPC cable of the device using a zero-insertion force (ZIF) connector (FH39 series, Hirose) on one side and the RHS 32-channel head-stage with the RHS2116 amplifier head-stage on the other side using an Omnetics connector (NPD-36-AA-GS, Omnetics). The amplifier head-stage was subsequently connected to the Intan RHS recording/stim system using a serial peripheral interface (SPI) cable. The device was designed to encompass an internal reference and ground which was reflected in the PCB layout. The sampling rate of the recording was 30 kHz, and it was performed in a grounded setup in an incubator which acted as a Faraday cage. An Arduino MEGA was also connected to the Intan RHS recording/stim setup and was used to trigger recordings every hour for a duration of 5 minutes.

### Data Pre-processing

#### Data Loading

For single file analysis, MEA data (.rhs and .rhd), was loaded onto NeuroMaps to extract the embedded information such as the recorded signals, timestamps, channel identifiers, impedance values (optional), and phase information (optional). For batch analysis, the data saved with identifiers such as ‘experimental ID’, ‘spikeData’, ‘lfpData’ was loaded into the ‘aggregate_analysis.m’ NeuroMaps module for longitudinal data analysis.

#### Quality Assessment

Electrodes were first screened based on impedance measurements, and any channel with impedance exceeding 1MΩ was excluded to eliminate non-functional or unstable electrodes. In the case where impedance measurements was not collected, this step was skipped. Following this hardware-level filtering, additional statistical and spectral metrics were used to identify noisy or artefactual channels. Channels were flagged if their standard deviation (STD) or median absolute deviation (MAD) exceeded 3.5 times the median across all channels, indicating abnormally high variance. Power spectral density (PSD) was then computed using Welch’s method to quantify frequency-domain characteristics of the signal. High-frequency noise was evaluated by measuring the power above 80% of the Nyquist frequency. Channels exceeding a high-frequency PSD threshold of 0.02 µV^2^/Hz were further assessed for coherence with the rest of the array. Any channel failing one or more of these criteria was excluded from downstream spike detection and analysis. This approach is based on methods developed by the International Brain Laboratory (IBL)^72^ and previously applied by Spikeinterface^42^.

#### Referencing and Filtering

The data of the corresponding good channels was filtered for powerline hum using an infinite impulse response (IIR) filter of order 4 between 50 Hz and 6 kHz at 100 Hz increments (harmonics) with a 2 Hz bandwidth around each harmonic. The cleaned data was subsequently referenced using a common median reference (CMR) rather than a common average reference (CAR). This method is of preference to avoid data biasing during high spiking activity^6^. Referenced data was then band-pass filtered (3^rd^ order Butterworth filter, 100-6000 Hz) to detect multi-unit activity. For low-frequency analysis, the powerline filtered data was down-sampled to 1 kHz and low pass filtered using a bi-directional finite impulse response filter (FIR) using *eegfilt.m* (EEGLAB)^73^. This was done at 500 Hz and at the sub-band frequencies previously reported^6^: delta (0.5-4 Hz), theta (4-7 Hz), alpha (8-13 Hz), beta (13-30 Hz), gamma (30-50 Hz) and broadband high gamma (100-400 Hz). Similar to previous work, high gamma analysis was performed in separate segments, 100-200 Hz and 200-400 Hz, to prevent biasing the power estimates towards lower frequency limits due to the inverse power law (1/f) that the power spectrum follows.

### Multi-unit Activity (LFP) Analysis

#### Spike Detection

Spikes were detected for each channel using a 60-second window and a threshold-based method, in which the threshold was set to 4.5σ_*n*_. This value can be adjusted based on the noise levels of the signal visualised in NeuroMaps. The noise level was robustly estimated using the formula, σ_*n*_= *median*(|*S*_*t*_|/0.6745), where |*S*_*t*_| is the absolute signal amplitude^43^. For spike detection, ‘findpeaks.m’ was used where a peak was defined by its minimum peak height (4.5σ_*n*_) and a minimum peak distance (3 ms). A maximum peak height was also set (30σ_*n*_) to eliminate spurious charges. Duplicate spikes were removed by sorting the spike locations of the concatenated positive and negative spikes and removing the lower amplitude spike occurring within the refractory period (2 ms). Non-physiological waveforms were also detected by identifying and eliminating instances where more than 3 spikes were detected within the refractory period. Moreover, spikes exceeding a full width at half-maximum (FWHM) threshold of 1 ms were considered non-biological and were excluded from further analysis. The spikes were centred around a 4 ms window.

#### Spike Sorting and Clustering

Preliminary clustering was performed as a general quality check of waveform separability. Waveforms were first z-score normalised, reduced in dimensionality using principal component analysis (PCA), and clustered using k-means with a minimum cluster size of 10. Mean and standard deviation waveforms were visualised for each cluster, and non-physiological events were discarded.

For more refined sorting, we applied a previously published method^74^ integrating wavelet decomposition with Gaussian mixture modelling (GMM). Briefly, waveforms were decomposed using a Haar wavelet transform, and the resulting coefficients were z-score normalised. Each wavelet component was weighted by a GMM-based separability metric prior to PCA. The top five principal components were retained, and up to 12 Gaussians were used to overfit the data in high-dimensional space. Final cluster centroids were refined using Nelder-Mead^75^ optimisation to identify local density maxima in feature space, and distinct spike clusters were accordingly labelled.

Following clustering, spike timestamps were used to generate a binary vector matching the length of the original signal, with ‘1’ indicating a spike event and ‘0’ elsewhere. This binary array was used to generate raster plot and in downstream population activity analyses.

#### Network Analysis

Population network activity was visualised by summing the binary spike arrays across all electrodes in 1 ms bins. The resulting population firing rate was smoothed using a 100 ms Gaussian kernel to reveal underlying network dynamics. Bin size and smoothing window were selected based on previously reported methods and parameters, to preserve temporal resolution while reducing noise^6,29^. Functional connections were identified using spike-time-tiling-correlation matrix (STTC)^45^ and the nodes were classified as receiver, sender, or broker nodes by computing the latency in spikes detected at each electrode and comparing the incoming and outgoing number of spikes such that they exceed 80%.

#### Spike Features and Synchronicity

Various features including full-width half maxima, peak-to-peak amplitude, inter-spike-interval (ISI), and bursts. Bursts were detected if they had a minimum duration of 50 ms and contained a minimum of 3 spikes with a maximum ISI of 100 ms. The synchronicity was evaluated in the case where more than 8 channels were active and computed using SpikeContrast^76^.

### Local-field Potential (LFP) Analysis

#### FOOOF Analysis and Oscillatory Power

To quantify oscillatory activity independently from aperiodic (1/f) components, power spectral densities (PSDs) were computed using Welch’s method (2 s window with 50% overlap). Prior to using the Fitting Oscillations and One-Over-F (FOOOF) toolbox^30^, the PSDs were interpolated at frequencies affected by powerline notch filtering (50 Hz and its harmonics) to smooth the dips which might confound the results. Subsequently, log power spectra were fit between 1 and 100 Hz. The FOOOF model separates the power spectrum into an aperiodic component (modelled as a linear function in log-log space) and a set of periodic peaks.

The aperiodic exponent and offset were extracted as measures of broadband activity. Oscillatory power was assessed by summing the power of identified peaks after subtracting the aperiodic fit. Only peaks with power above a defined threshold and goodness-of-fit exceeding R^2^>0.9 were retained. Parameters for FOOOF were based on previous work^30^.

#### Phase Amplitude Coupling (PAC) and Modulation Index (MI)

For each of the filtered sub-bands, the instantaneous phase and amplitude were extracted using a Hilbert transform (Hilbert.m, angle.m) while the power was extracted by squaring the amplitude of the filtered sub-band signal. Phase amplitude coupling (PAC) was computed by taking the instantaneous sub-band phases and binning them into 20 equidistant bins between −π to π. The corresponding power was sorted based on the corresponding phase and visualised by circularly shifting it such that the maximum power was at −π. The modulation index (MI) was used to quantify the phase-amplitude coupling strength. This measure was derived from the Kullback-Leibler (KL) distance^34^, which quantifies the divergence of the sub-band’s power compared to a uniform distribution, across the phase bins.

#### Continuous Wavelet Transform (CWT)

Continuous wavelet transform was used to decompose the low-pass filtered neuronal signal into specific frequency bands to obtain time-frequency signal representations while preserving both spectral and temporal information.

### Functionality testing

The brain organoids were recorded in BrainPhys medium. For functionality testing, 50 mM potassium chloride (KCl) was used to modify the electrical activity of the organoids. The baseline measurements were obtained before and after the addition of the reagent. The sample was rinsed three times with Dulbecco’s PBS (DPBS) and a fresh medium was added after each reagent treatment.

